# Standardization of suspension and imaging mass cytometry readouts for clinical decision making

**DOI:** 10.1101/2023.03.19.531228

**Authors:** Ruben Casanova, Shuhan Xu, Sujana Sivapatham, Andrea Jacobs, Stefanie Engler, Tumor Profiler Consortium, Mitchell P. Levesque, Reinhard Dummer, Bernd Bodenmiller, Stéphane Chevrier

## Abstract

Suspension and imaging mass cytometry are single-cell, proteomic-based methods used to characterize tissue composition and structure. Data assessing the consistency of these methods over an extended period of time are still sparse and are needed if mass cytometry-based methods are to be used clinically. Here, we present experimental and computational pipelines developed within the Tumor Profiler clinical study, an observational clinical trial assessing the relevance of cutting-edge technologies in guiding treatment decisions for advanced cancer patients. By using aliquots of frozen antibody panels, batch effects between independent experiments performed within a time frame of one year were minimized. The inclusion of well-characterized reference samples allowed us to assess and correct for batch effects. A systematic evaluation of a test tumor sample analyzed in each run showed that our batch correction approach consistently reduced signal variations. We provide an exemplary analysis of a representative patient sample including an overview of data provided to clinicians and potential treatment suggestions. This study demonstrates that standardized suspension and imaging mass cytometry measurements generate robust data that meet clinical requirements for reproducibility and provide oncologists with valuable insights on the biology of patient tumors.

## Introduction

Cytometry by time of flight (CyTOF), also known as suspension mass cytometry, as well as imaging mass cytometry (IMC) are technologies capable of simultaneously measuring more than 40 parameters at the single-cell level; the data generated by these mass spectroscopy-based methods are consistent with flow cytometry and immunohistochemistry measurements (1–5). High-dimensional characterization of complex tissues at single-cell resolution has improved our understanding of immunology and tumor biology (6–10). The democratization of single-cell technologies has led to large-scale projects aimed at characterizing human tissue at single-cell resolution with the goal of providing valuable resources for clinicians (11). For example, the comprehensive characterization of immune populations or molecular signatures of tumor tissue can provide insight into the most appropriate therapeutic approach, as highlighted by clinical trials that employ mass cytometry to evaluate the response to immunotherapy in patients with endometrial, non-Hodgkin B cell lymphoma, and breast cancer (12). Transitioning a technology from the academic research setting to clinical laboratories requires standardized protocols, proper quality control protocols, and robust experimental and analytical pipelines (13–15). Additionally, for clinical translation, different facilities must be able to use the technology, and turnaround times for measurement and analyses must be short. Such a transition from a research setting to a clinical setting therefore requires optimization of many methodological factors in order to achieve measurements reproducibility and scalability.

Several approaches have been proposed to minimize batch effects across mass cytometry measurements including the use of frozen or lyophilized antibody cocktails, shared references across independent experiments, and a fully standardized data analysis pipeline (16–21). Although this type of efforts has assessed data consistency across institutions of an immunofluorescence imaging workflow based on the Opal Vectra system (22), no reports assessing data consistency of highly multiplexed, single-cell cytometric platforms over time or sites are available.

In the context of the Tumor Profiler (TuPro) study, an observational clinical trial involving collaborative work at the ETH Zurich, the University of Zurich, the University Hospital Zurich, the University Hospital Basel, Kantonsspital Basel, and Roche Holding AG (23), we developed experimental and computational pipelines to standardize CyTOF and IMC readouts with the aim of rendering these technologies clinically applicable. Through TuPro, over a period of two years, 240 samples from patients with melanoma, ovarian cancer, or acute myeloid leukemia were analyzed on exploratory platforms including CyTOF, IMC, single-cell RNA and DNA sequencing, bulk RNA sequencing, proteotyping, and ex vivo drug testing. TuPro aimed to assess the technical feasibility of these platforms to deliver data within a two-week turnaround time with sufficient throughput.

TuPro data and treatment suggestions based on these data were provided to clinical teams as were routine pathology and FoundationOne® test data. For each treatment suggestion, the contributions of individual technologies were recorded. This setup allowed for a precise evaluation of the impact of each technology on clinical decisions, which highlighted the high value provided by both CyTOF and IMC to the clinic (manuscript in preparation). Here, we describe the experimental and computational pipelines developed in the context of the TuPro trial that could be applied more broadly to inform treatment selection. This report provides our evaluation of the consistency of the data generated using CyTOF and IMC on the melanoma cohort and shows the readouts returned to the clinician for a representative case study.

## Results

### Analytical pipelines for CyTOF and IMC to ensure reproducibility and stability

The sample processing pipelines for CyTOF and IMC were tailored to the needs of the TuPro trial. This observational clinical trial was designed to provide clinically relevant data within a time frame of two weeks while ensuring data consistency over two years (Figure 1A). For CyTOF, dissociated tumor material was barcoded (24), allowing the simultaneous analysis of up to five tumors and their matched peripheral blood mononuclear cells (PBMCs) in a single run.

**Figure 1.**
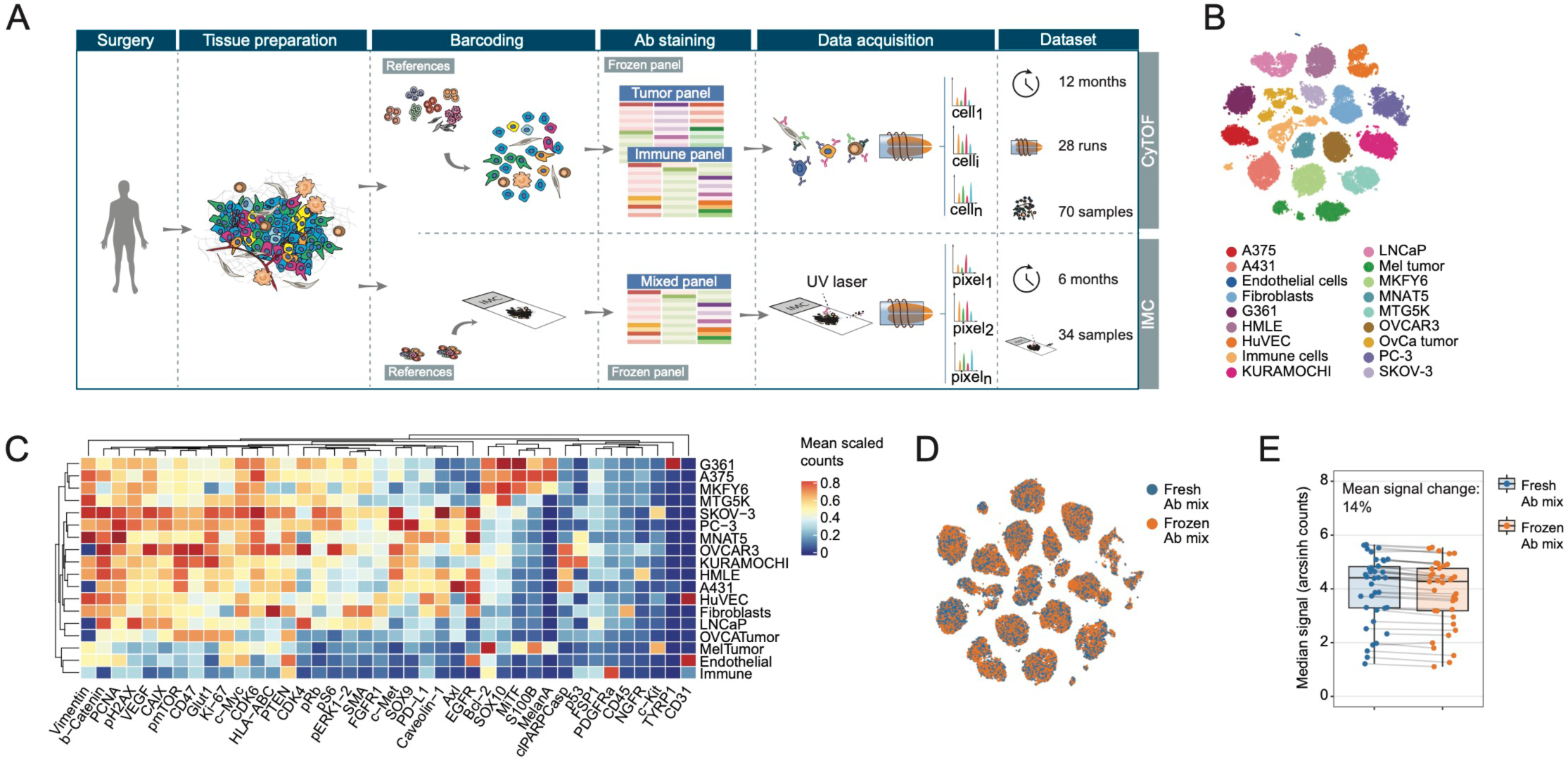
Standardized CyTOF and IMC workflows to assess clinical samples based on well-defined anchoring reference sets and frozen antibody panels. A. Schematic view of the workflow optimized for the TuPro observational clinical trial. B. t-SNE representations of a maximum of 3,000 cells from each of the 18 reference samples used as positive and negative controls for the CyTOF tumor panel. C. Heatmap of the mean scaled count of each of the markers in the CyTOF tumor panel in each of the reference samples used for CyTOF. D. t-SNE representations of a maximum of 3,000 cells from each reference samples stained with the fresh and the frozen versions of the CyTOF tumor panel. Cells on the t-SNE are colored by antibody panel type. E. Dot plot showing the median signal intensity for the reference with the highest expression level for each marker upon staining with the fresh and frozen CyTOF tumor panel. The markers in each pair for fresh vs. frozen panels are connected with a line.

In order to provide clinicians with an in-depth characterization of both the immune and the tumor compartments, CyTOF analysis was performed with two antibodies panels: an immune panel of 38 antibodies to distinguish immune cells, including B cells, T cells, granulocytes, monocytes, macrophages, and plasma cells, and a tumor panel of 41 antibodies to tumor markers enabling characterization of diverse tumor phenotypes. We monitored signal stability over 1 year. For IMC, a single antibody panel with 38 antibodies was used to simultaneously characterize a variety of immune cells as well as tumor cells in 34 samples over 6 months.

Each analytical run included well-characterized cell lines as positive and negative controls for each marker in the panel to assess data consistency throughout the trial (Figure 1B-C and Figure S1A-D). For IMC, a tissue microarray assembled from cell pellets from a selection of Melanoma, Ovarian cancer cell lines, PMBCs, and a sample of tonsil tissue, was used as reference. For CyTOF, the reference samples were included with the samples upon barcoding, whereas for IMC, references were loaded onto the histological slide next to the tumor section (Figure 1A).

In order to minimize antibody concentration variability across independent experiments, a frozen antibody mix was generated at the beginning of the study. This ensured that identical aliquots were used throughout the trial. We compared the staining profile of each antibody between fresh and frozen panels and observed no obvious differences in the t-SNE visualization or in the density plots for either the IMC panel or the CyTOF panels (Figure 1D, Figure S1E-G); this is consistent with previous work showing that antibody freezing does not affect staining pattern significantly (20). A more precise quantification of the signal intensity obtained upon staining with the fresh versus the frozen antibody panels revealed an average loss of 13% (range: -26% to +8%) for the antibodies included in the CyTOF tumor panel, of 4% (range: -35% to + 31%) for the antibodies included in the CyTOF immune panel, and of 11% (range: -27% to +8%) for the antibodies included in the IMC panel (Figure 1E, Figure S1H-I). Together, these data confirmed that using a frozen antibody mix does not lead to significant changes in staining efficiency, and that a reference set provides relevant positive and negative controls that allow data consistency to be assessed over an extended period of time.

### Data stability achieved over one year of acquisition

We used the reference set to assess data consistency in 28 CyTOF runs performed over 12 months and 34 independent IMC experiments performed over 6 months. For CyTOF, we observed a large variation in the number of cells acquired, which could be partly explained by the differences in numbers of tumor and PBMC samples analyzed in the same run (Figure 2A). Despite the difference in absolute cell numbers, the frequency of the different cell types was stable across the runs with an average coefficient of variation (CV) of 14.2% (range: 7.3-29.5%, Figure 2B-C). In absolute numbers, the largest differences were observed between immune and non-immune cells, suggesting potential bias in the transmission efficiency based on cell size (Figure S2A). The average CV observed with the immune panel on the cell types identified based on a random forest classifier was of 17.7% (range: 6.4-46.2%, Figure S2B-C). For IMC, the numbers of cells of the different populations identified based on a random forest classifier after segmentation were more stable (CV=11.09%, Figure 2D), but frequencies were less stable (average CV=20.8%, range: 11.1-33.4%, Figure 2E-F), than with CyTOF. Endothelial cells were assessed in tonsils, with expected frequency variation from one section to another. Therefore, this cell type was excluded from the frequency analysis and included only for marker expression assessment.

**Figure 2.**
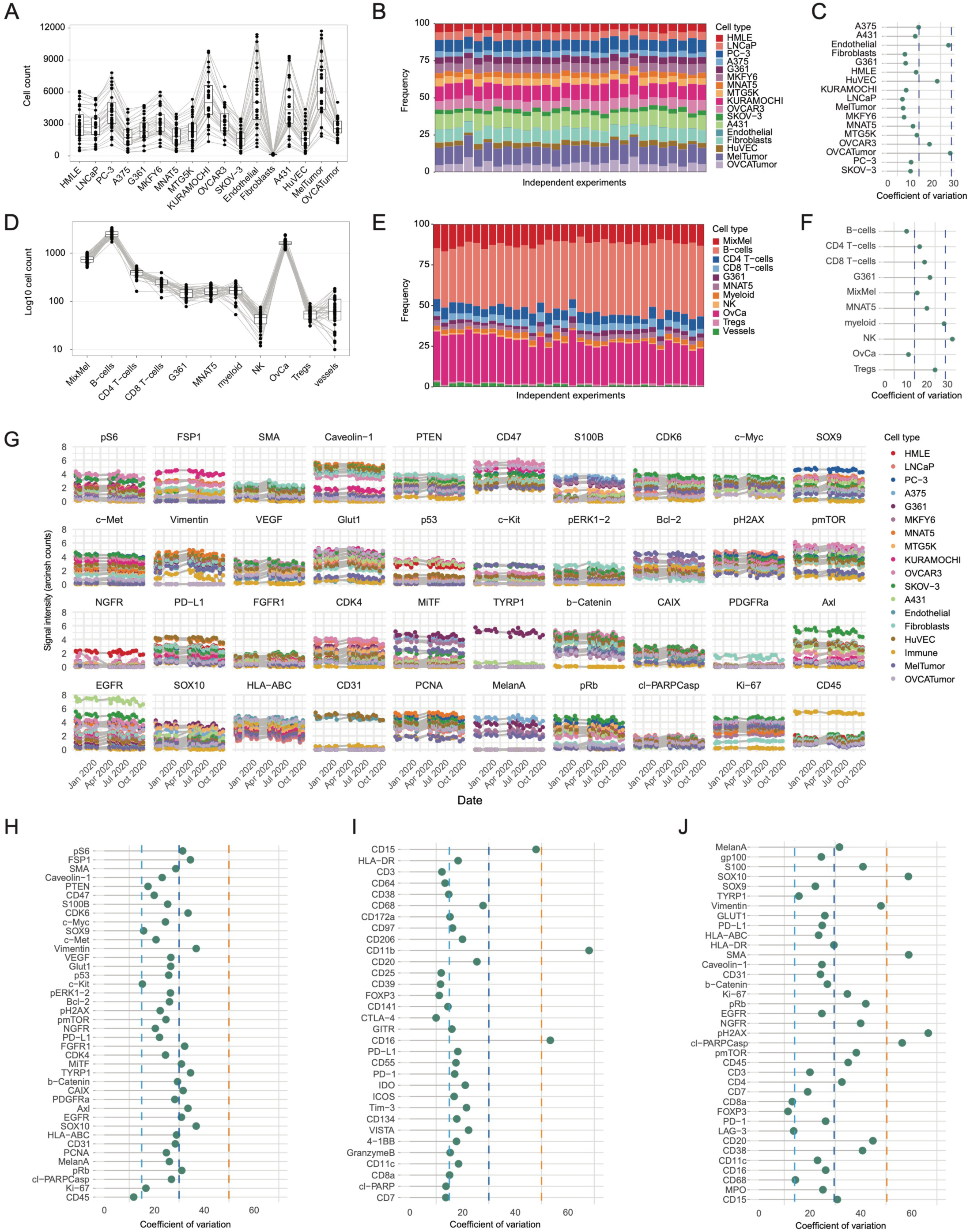
Data stability achieved over a year of acquisition. A. Box plot of the numbers of cells of particular types identified in reference samples based on a random forest classifier across all CyTOF experiments using the tumor panel. Cell types collected in the same acquisition are connected with lines. B. Stacked bar plot of cell-type compositions of reference samples identified in each experiment acquired with the CyTOF tumor panel. C. Dot plot of the CVs of the frequencies of cell types identified reference samples acquired with the CyTOF tumor panel. Dashed lines indicate CVs of 15% and 30%. D. Box plot of the numbers of cells of particular types identified in reference samples based on a random forest classifier across all IMC experiments. Cell types collected in the same acquisition are connected with lines. E. Stacked bar plot of cell-type compositions of reference samples identified in IMC experiments. F. Dot plot of CVs of the frequencies of cell types identified in reference samples by IMC experiments. Dashed lines indicate CVs of 15% and 30%. G. Dot plots of median signal intensities of indicated markers in reference samples across all measured time points for data acquired with the tumor CyTOF panel. Marker expressions in each cell type across experiments are connected with lines. H-J. Dot plots of CVs of median signal intensities of indicated markers in reference samples for each marker for H) data acquired with CyTOF using the tumor panel, I) data acquired with CyTOF using the immune panel, and J) with IMC.

The signal stability of each marker was assessed based on the median expression level in cell lines with the highest signal, after exclusion of cell subsets with a frequency below 1% (Figure 2G). The CV for the tumor markers assessed using CyTOF showed limited variation over several months with an average CV of 26.4% (range: 11.8-36.9%, Figure 2H). The CV of immune markers was in the same range with an average of 20.5% (range: 10.0-68.0%, Figure 2I). We observed three outliers in this analysis, CD11b, CD15, and CD16; these markers were assessed on granulocytes, which displayed a high variability across batches for most markers assessed for reasons which are difficult to apprehend (Figure S2D-F). IMC marker variation was slightly higher with an average CV of 31% (range: 11.5-66.6%, Figure 2J and Figure S2G). Together, these data demonstrated that our standardized pipeline led to the generation of stable data for both IMC and CyTOF over extended periods of time.

### Linear scaling using references allows for efficient batch correction

In order to correct for the signal variation identified between runs (Figure 2G and Figure S2D, G), we carried out a batch correction using linear scaling for each individual marker based on a fixed percentile of the signal identified for the most positive reference cell line as previously described (19,25). A channel-specific factor was calculated by dividing the 50^th^ percentile assessed in a specific run by the average 50^th^ percentile calculated across all experiments (Figure 3A). The signal in each channel was then divided by this correction factor, which aligned the distribution to the selected percentile (Figure 3B, Figure S3A-C). A systematic assessment based on principal component analysis performed on all cells from the relevant reference and taking into account all channels, revealed that the distributions of cells in the references were more compact (larger average silhouette coefficient) after batch correction compared to the same analysis performed on raw data (Figure 3C). We made similar observations on the immune CyTOF panel and the IMC panel (Figure 3D-E), confirming that this method efficiently corrected for batch-to-batch variation.

**Figure 3.**
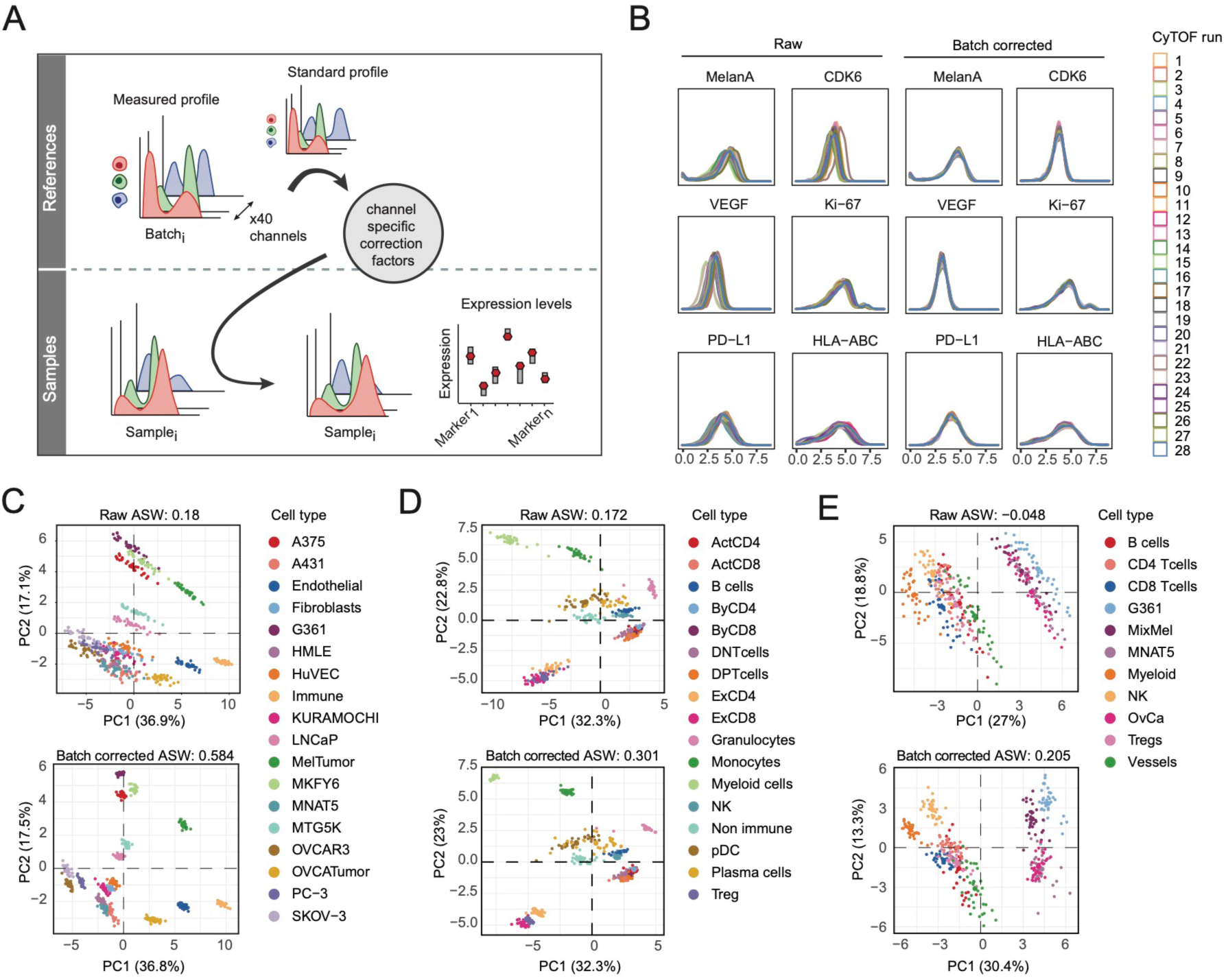
Batch correction upon linear scaling based on a percentile. A. Overview of the batch correction approach for CyTOF. For each new acquisition, the median signal intensity of each marker is calculated on reference sample with the highest level of expression for that marker (top left) and a channel specific correction factor is derived (top right). The derived correction factors are applied to the measured samples (bottom right) in order to retrieve corrected staining intensities (bottom left). B. Density plots for selected markers included in the tumor panel before (left) and after (right) batch correction. C-E. Principal component analysis projections of the acquisitions before and after batch correction colored by reference sample for C) the CyTOF tumor panel, D) the CyTOF immune panel, and E) the IMC panel. The average silhouette widths (ASWs) are indicated for raw and batch corrected data.

### Validation of batch correction on independent tumor samples

In order to assess the effect of batch correction on patient samples, we took advantage of a pooled melanoma sample prepared by combining samples from three individuals. This pooled sample was evaluated in each CyTOF run in the trial to serve as an internal control for assessment of batch-to-batch normalization. In the absence of normalization, the main cell types identified in this sample, which included immune, tumor, endothelial, and fibroblast cell types, were well aligned on the t-SNE map (Figure 4A). Cell proportions were stable across this study, particularly for abundant cell types, with CVs of 7.5% for tumor cells and 11% for immune cells (Figure 4B, C). CVs of endothelial cells and fibroblasts were in the range of 20%, whereas apoptotic cells and undefined cells had higher variation (Figure 4C). Similar observations were made on the tumor compartment. No batch effects were detected when tumor cell phenotypes were identified by unsupervised clustering using FlowSOM (Figure 4D-E). For all clusters with a frequency above 1%, the CVs were below 30%.

**Figure 4.**
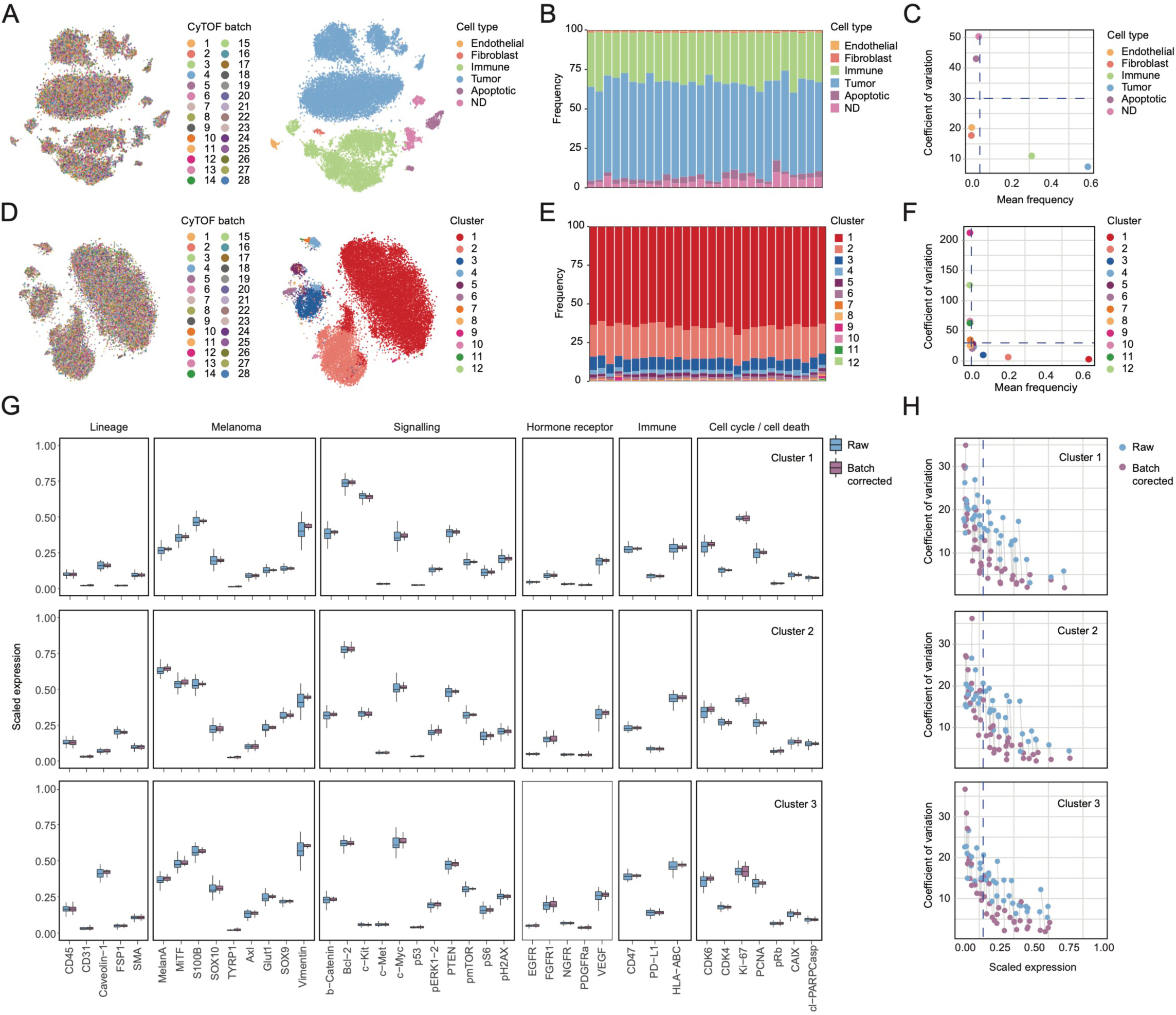
Validation of batch correction in an independent tumor sample. A. t-SNE representation of a maximum of 1,000 cells from the validation tumor sample for each acquisition batch colored by batch (left) and cell type (right). B. Stacked bar plot of cell-type frequencies identified in each acquisition with the CyTOF tumor panel. C. Dot plot of the relationship between the CVs and mean cell-type frequencies. Horizontal line indicates 30% CV; vertical line indicates 5% frequency. D. t-SNE representation of a maximum of 1,000 tumor cells from the validation tumor sample for each acquisition batch colored by batch (left) and cell type (right). E. Stacked bar plot of tumor cluster frequencies identified by unsupervised clustering for each acquisition with the CyTOF tumor panel. F. Dot plot of the relationships between the CVs and mean cluster frequencies. Horizontal line indicates 30% CV; vertical line indicates 1% frequency. G. Boxplots of raw and corrected scaled marker intensities across all acquisitions for the three most abundant tumor clusters. H. Dot plots of relationships between the CVs and scaled marker expression (raw vs. batch corrected) for the three most abundant tumor clusters. Each pair of markers (raw vs. batch corrected) is connected with a line. Vertical line indicates 0.15 mean scaled intensity.

A key readout provided to the clinicians was the marker expression in the main tumor subpopulations. In order to determine the accuracy of this readout, the mean expression of each marker included in the CyTOF tumor panel was assessed in the three main tumor subsets (C1, C2, and C3) before and after signal correction (Figure 4G). The highest CVs were around 30% before batch correction, and CVs decreased with increasing signal intensities. Batch correction led to a significant decrease in median CV (15.7% before correction to 8.0% after correction) across the measured markers for three main tumor subsets (Figure 4H and Figure S4B-C). As IMC-based tissue imaging cannot be performed multiple times on the same tumor sample, such an approach could not be performed for evaluation of the effect of batch correction on IMC data. Analyses of CyTOF data demonstrated, however, that the CVs of the reported signal intensities were systematically below 10% when the 0 to 1 normalized expressions were above 0.2.

### Single-cell readouts for targeted treatment decision support

Single-cell readouts focusing on tumor heterogeneity as well as immuno-scores were presented to molecular tumor boards for clinical decision support in the TuPro trial. We illustrate this here for a 61-year-old male, diagnosed with stage IV cutaneous melanoma cancer, who progressed despite anti-PD-1 checkpoint inhibitor treatment. TuPro technologies were used to analyze a biopsy of an inguinal lymph node metastasis from this patient. The CyTOF data were based on the analysis of approximately 10,000 live cells with both immune and tumor panels, whereas IMC analysis was based on 40,000 segmented cells (Figure S5A). The sample was dominated by immune cells (>50%) and contained 25% of tumor cells (Figure 5A). A few percent of cells were identified as endothelial cells, and a fairly large percentage of the remaining cells expressed DNA damage markers.

**Figure 5.**
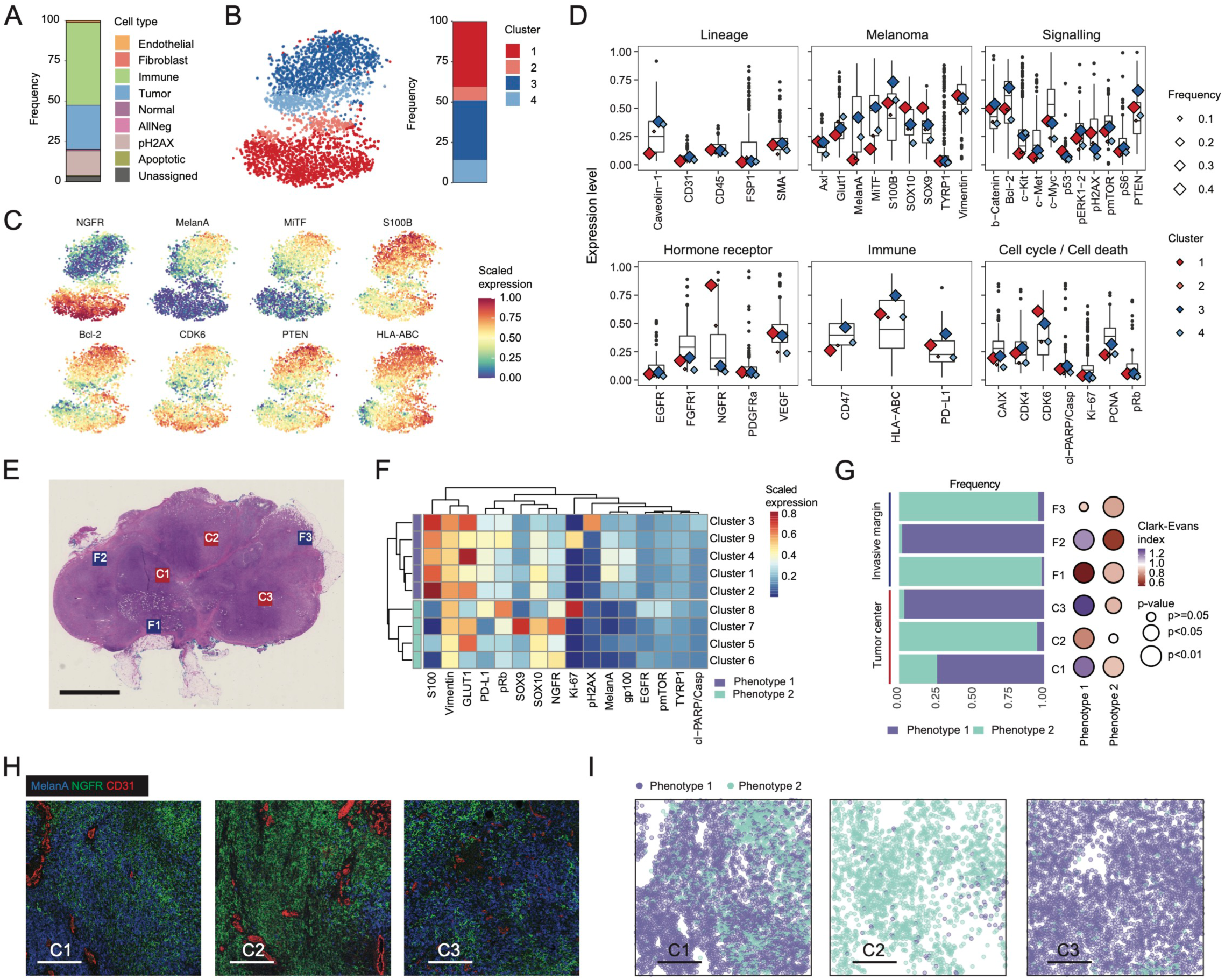
Single-cell readouts of the tumor compartment of a metastatic melanoma cancer patient for clinical decision support. A. Stacked bar of cell-type frequencies of a sample from a metastatic melanoma cancer patient based on CyTOF analysis. B. t-SNE representation of a maximum of 1,000 tumor cells colored by FlowSOM cluster. C. The same t-SNE representation colored by scaled intensities from a selection of melanoma markers. D. Boxplots of distributions of intensities of melanoma markers on tumor cells across the TuPro cohort. Rhombuses indicate the mean intensities for a given cluster. Sizes and colors correspond to frequencies and cluster identity, respectively. E. Hematoxylin and eosin-stained FFPE section corresponding to the sample analyzed by IMC. Regions selected for IMC analysis are indicated with red and blue boxes for central and peripheral regions, respectively. Scale bar, 5 mm. F. Heatmap of mean scaled marker intensities in each identified FlowSOM cluster, subsequently grouped into two main phenotypes based on hierarchical clustering using Euclidean distance and Ward.D2 linkage. G. Stacked bar plot of frequencies of identified phenotypes split by imaged regions (left) and dot plot of Clark-Evans scores of the same region with color scale and dot size correspond to scores and p-values, respectively (right). H. IMC images showing expression of MelanA (blue) NGFR (green) and CD31 (red) in the three central regions of the patient sample. Scale bar, 250 µm. I. IMC images of the regions shown in H with each cell identified as a dot colored by tumor phenotypes. Scale bar, 250 µm.

In-depth analysis of the tumor compartment based on markers specific for melanoma subtypes, signaling pathways, hormone receptors, immune ligands, and markers of cell-cycle stage revealed the presence of two main clusters (C1 and C3) characterized by different levels of NGFR, MelanA, and MiTF (Figure 5B-C). The expression of each marker was compared to the median expression level found in the first 24 samples measured during the TuPro study. Cluster C1 identified in this patient was among the highest in terms of NGFR expression, whereas cluster C3 was in the high range for MelanA, MiTF, and S100B (Figure 5D). Cells in both clusters expressed high levels of CDK6 and of the immune-related markers HLA-ABC and PD-L1 compared to the median of the background melanoma cohort. Most tumor cell clusters also expressed high levels of PTEN and ß-catenin but low levels of c-Met and c-Myc. To provide more insight regarding the spatial distribution of the different cell types, IMC data were generated on six regions of interest located at the center of the tumor and at the invasive margin (Figure 5E). The analysis of the tumor compartment confirmed cell plasticity with the presence of two main melanoma cell phenotypes driven by their mutually exclusive expression of MelanA, S100, and NGFR; these cells heterogeneously expressed other tumor markers such as GLUT1 and PD-L1 (Figure 5F, Figure S5B-C). The presence of tumor cells that express high levels of NGFR have been linked to resistance development during immunotherapy (26). The main tumor phenotypes identified using unsupervised clustering of IMC data showed some degree of spatial compartmentalization in both central and peripheral regions (Figure 5G left, Figure S5D-E). We computed a simple measure of spatial aggregation among tumor cells for each image using the Clark-Evans index, which is the ratio of the mean distance to the nearest neighbor in a point pattern to the expected mean distance if that pattern was randomly distributed (27). This analysis showed different spatial aggregation patterns between the two identified tumor phenotypes with a moderate, yet consistent spatial clustering for phenotype 2, in contrast to phenotype 1, which showed spatial spreading in three of the six imaged regions (Figure 5G, right). All detected tumor clusters showed various degrees of spatial aggregation with Clark-Evans scores particularly low for GLUT1^+^ tumor cells (clusters 4, 5, and 7), indicating a strong spatial aggregation of these cells in response to hypoxia (Figure S5F).

CyTOF analysis of the immune landscape of the melanoma sample revealed the presence of about 50% T cells and 40% myeloid cells (Figure 6A, B). The CD8/CD4 ratio was close to 2, among the highest observed in the background melanoma cohort (Figure 6C). The CD8^+^ T cell compartment was dominated by exhausted T cells, whereas the CD4^+^ compartment was dominated by regulatory T cells (Tregs) (Figure 6D). Exhausted T cells were characterized by high levels of CD38, CD39, Tim-3, CD27, ICOS, Granzyme B, and TIGIT and by low levels of PD-1 (Figure 6E, Figure S6A). In exhausted T cells from patients who have been treated with anti-PD-1 immunotherapy, as was this patient, we usually detected low levels of PD-1 by CyTOF (manuscript in preparation). This effect may be due to competitive binding between the anti-PD1 used for treatment and the PD-1 antibody clone used in CyTOF, since PD-1 was observed in immune cells by IMC analysis with a different antibody clone (Figure S6B). Myeloid cells were characterized by high levels of immunoregulatory markers such as CD38, PD-L1, IDO, and TIM3 (Figure 6F).

**Figure 6.**
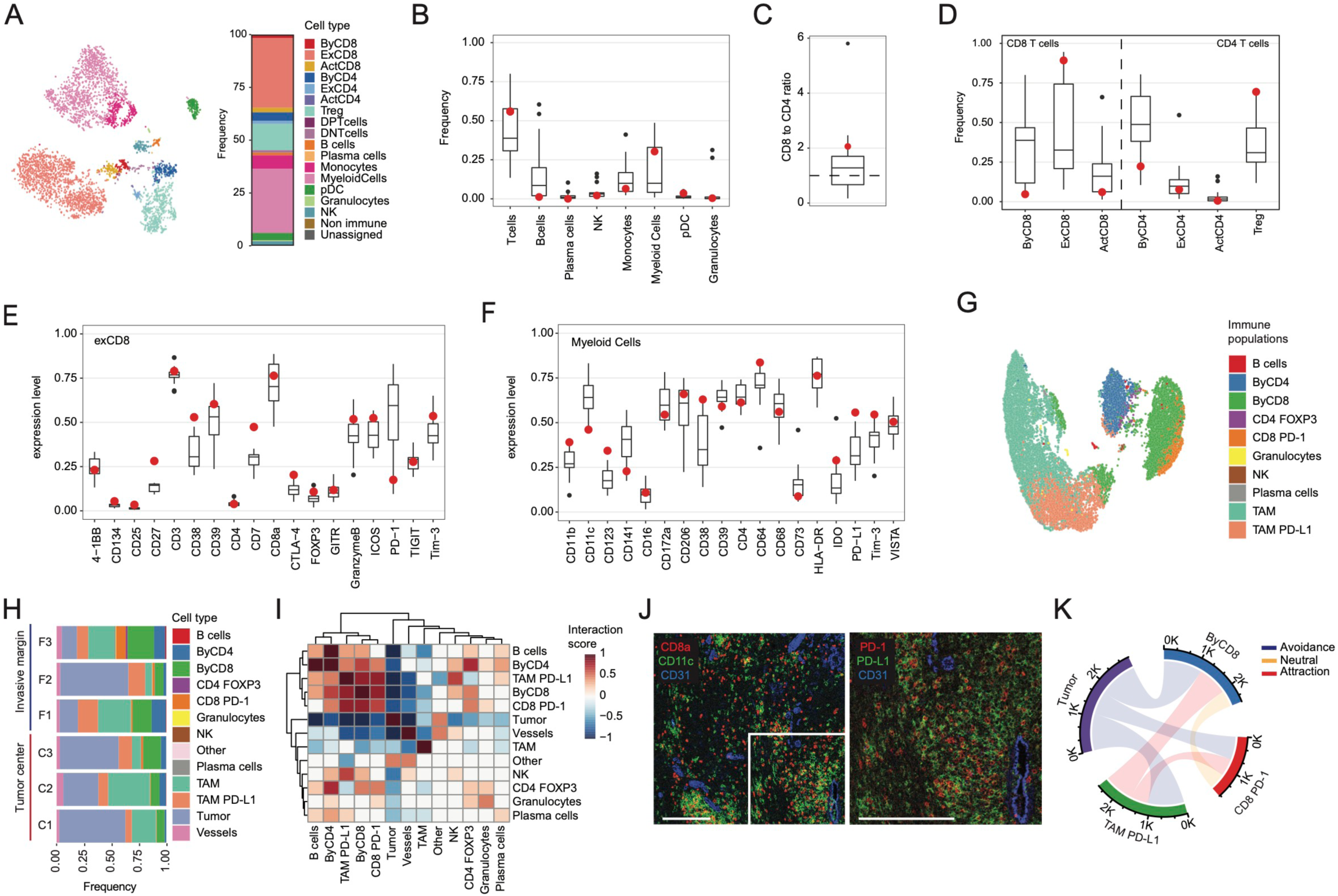
Single-cell readouts of the immune compartment of a metastatic melanoma cancer patient sample for clinical decision support. A. t-SNE representation of a maximum of 1,000 immune cells colored by cell type, identified using a random forest classifier (left), and corresponding cell-type frequencies shown on a stacked bar plot (right) of the sample from the metastatic melanoma cancer patient. B-D. Boxplots of the distributions of B) immune cell frequencies, C) CD8^+^ to CD4^+^ T cells ratios, and D) detailed T cell populations across the TuPro. Red dots indicate the values for the patient sample analyzed for this study. E-F. Boxplots of distributions of intensities of E) T cell markers and F) myeloid markers across the TuPro cohort. Red dots indicate the values for the sample analyzed for this study. G. UMAP representation of a maximum of 20,000 immune cells identified using a random forest classifier. H. Stacked bar plot of the frequencies of identified immune cells split by imaged regions. I. Mean interaction scores resulting from the neighborhood permutation test performed on all cell pairs for each imaged region. Scores range from -1, indicating avoidance, to +1, indicating cell co-localization. J. Representative IMC image showing CD8^+^ T cells in red, CD11c^+^ tumor-associated macrophages in green, and blood vessels in blue. Scale bar, 250 µm. K. Circular plot illustrating the interactions between tumor cells, PD-L1^+^ tumor-associated macrophages, CD8^+^ T cells (PD-1^+^ and PD1^-^) as well as tumor cells. Edge width corresponds to the total number of cells in contact with each other, and the color corresponds to the result of the permutation testing. Red: attraction, blue: avoidance, yellow: neutral.

IMC analysis confirmed high frequencies of both CD4^+^ and CD8^+^ T cells as well as macrophages in the images from this patient compared to those from the background melanoma cohort (Figure S6C). IMC also indicated relatively high proportions of PD-1^+^ CD8^+^ T cells and PD-L1^+^ macrophages (Figure 6G-H, Figure S6C). Analyses of detailed compositions of each region of interest revealed high frequencies of immune cells in both central and marginal regions (Figure 6H). In order to further characterize spatial interactions across the sample, a neighborhood permutation test described previously (28) was carried out for all possible cell-pairs across all IMC images (Figure 6I). The data indicate a spatial colocalization between PD-1^+^ CD8^+^ T cells and PD-L1^+^ macrophages and avoidance between tumor cells and immune cells. A representative IMC image taken in a central region of the tumor highlights a significant immune infiltration with a uniform distribution of CD8^+^ T cells across the sampled region and the spatial proximity of T cells expressing PD-1 and macrophages expressing PD-L1 (Figure 6J). This was confirmed by permutation analysis of the same region (Figure 6G).

Based on these observations, the resistance to immunotherapy in this patient cannot be explained by an immune-excluded phenotype or by the downregulation of HLA-ABC on the surface of tumor cells. The induced expression of HLA-ABC and immune infiltration could be due to inflammation resulting from the anti-PD-1 treatment. It is unknown if the subpopulation of tumor cells that express high levels of NGFR existed in the tumor prior to anti-PD1 treatment or if cells underwent rapid dedifferentiation induced by inflammation as described previously (29).

Upregulation of CDK6, a direct target of Cyclin D1, is one of the strongest tumor-intrinsic predictors of resistance to immunotherapy (30). The upregulation of CDK6 observed in this patient suggests that this pathway contributed to the lack of response to anti-PD-1. These data suggest that the use of CDK4/6 inhibitors, shown in preclinical mouse studies to enhance the efficacy of PD-1/PD-L1 inhibitors and currently in clinical testing as a combinatorial therapy with immune checkpoint blockade in multiple solid tumors, could be an attractive treatment option for this patient. In addition, based on our immune cell profiling, we suggested treatments based on novel checkpoint inhibitor combinations such as those including anti-CTLA-4, anti-Tim-3, or anti-TIGIT. Biomarkers predictive of response to these combinations are not available, limiting their use in clinical practice. For this case study, the in-depth characterization of the tumor microenvironment allowed to better understand the resistance mechanism at play in this patient and provided highly valuable information to guide the clinicians in their treatment decision process.

## Discussion

CyTOF and IMC are two single-cell proteomic-based approaches that allow for an unbiased quantification of tumor sample composition, including the immune cell populations. Robust experimental designs are key for future incorporation of these technologies into clinical trial research and patient care. This study presents the experimental and computational pipelines developed to fulfill the requirements of the TuPro observational clinical study, in particular regarding long term data consistency, fast turnaround time and actionable target identification. To optimize data consistency, we made use of frozen antibody panels and confirmed that this process did not alter the measured protein signal compared to fresh antibody cocktails. The assessment of the accuracy of our readouts over time was based on cell type frequency and marker expression profile of well-characterized cell lines as well as primary cells used as references. The cell compositions of the reference samples were stable for both CyTOF and IMC platforms, with CVs for most populations well below the 20% commonly adopted as acceptance criteria for clinical application of similar technologies (31–33). Due to expected variation in cell composition across tissue sections, cell frequency stability in IMC could only be assessed in sections obtained from homogenously reconstituted mixed cell pellets.

The marker expression analysis preformed on the reference cells in CyTOF and IMC did not lead to the identification of any obvious signal drift, suggesting that frozen antibody panels are stable for at least one year. In CyTOF, signal intensities were first corrected for instrument sensitivity using bead-based correction (34) and at a second step by applying a linear scaling based on a percentile calculated on references. Since the references were processed together with the samples of interest, this second step allows to correct not only for instrument sensitivity but also for signal variation due to the staining procedure. In addition, the correction factors were determined for each measurement channel independently, allowing for a more precise correction of the instrument sensitivity compared to the bead-based correction, which is established only on five channels. When looking at the accuracy of the marker expression provided as the final readouts to the clinicians after batch correction, the CVs for most markers were again well below the 20% threshold commonly used as an acceptance criterion for clinical use. Conversely to CyTOF, IMC readouts cannot be normalized for instrument sensitivity based on bead normalization, which explains in part why the CV calculated on raw data tended to be higher for IMC compared to CyTOF. In addition, the assessment of the accuracy of the final readout returned to clinicians was prevented by the fact that homogenous tumor material, which can be repeatably measured over extended period of time cannot be obtained from consecutive sections of tumor tissue due to the inherent heterogeneity of the sample. The fact that signal detection in CyTOF and IMC are based on two modalities of the same instrument suggests that a similar level of data accuracy can be obtained for both methods.

The standardization of the experimental and computational pipelines developed to increase data consistency also led to decreased turnaround time. Throughout the study, both CyTOF and IMC systematically returned data for the samples provided within the two-week turnaround time defined at the beginning of the study. In addition, both methods demonstrated their ability to provide reports in as little as 72h post sample reception.

In term of actionable target identification, CyTOF and IMC have different properties, which make them highly relevant and complementary to provide added value to the clinic. CyTOF is a robust method for cell composition and phenotypic analysis that provides readouts derived from intact single cells, which can easily be distinguished from debris and doublets and the data collection can be readily standardized to integrate internal controls including beads and reference cells to ensure high data consistency. CyTOF is particularly well suited for fast acquisition runs, thus reducing instrument-related costs. In fact, the measurement of 50,000 cells could be performed in around 10 minutes at 100 cells/second, which was the introduction rate used in the TuPro study to minimize doublet generation. This contrasts with the more costly and time-consuming single cell technologies such as single cell RNA or DNA sequencing. As for those methods, CyTOF requires the use of fresh tissues, which necessitates adapting the logistics with hospitals to allow the measurement of single cells derived directly from liquid biopsies such as ascites or PBMCs, or after dissociation of viable solid tumor samples. For the latter, viable tumor material is typically not available for routine diagnostic purposes and needle biopsies typically do not yield enough material for CyTOF analysis, especially since the tissue dissociation steps will lead to sample loss (35).

For both of those aspects, IMC comes with obvious advantages. Since the analysis is done on the same type of tissue sections used in routine pathology, the IMC workflow is fully compatible with the clinic. In addition, the sample is analyzed in its native form and no bias is introduced due to sample preparation. The analysis can be focused on regions of interest selected based on expert knowledge of highly trained pathologists, and if additional information is required by the clinicians, the IMC setup also allows for follow up investigations based on the analysis of additional regions of interest present on the same tissue section, or on subsequent tissue sections. All the readouts generated include spatial cell localization, allowing to go beyond cell composition and phenotyping to also take into account the spatial proximity between cell types. Therefore, IMC allows to map cells within different compartments of the tumor tissue, which is particularly suited to score immune infiltration patterns. In the context of the TuPro study, six regions (three central and three peripheral) per sample with dimensions of 1 mm^2^ each were selected to maximize the likelihood of detecting all cell phenotypes present in the tissue (36). In terms of throughput, collecting the average 50,000 cells per sample took around 8 hours for IMC with a laser ablation speed of 200 Hz, and this throughput will considerably increase with the release of the next generation of the instrument.

The longitudinal study described here demonstrated the signal stability and reproducibility of CyTOF and IMC multidimensional data and showed that both methods provide an accurate and reproducible quantification of the tumor immune microenvironment over several months during the final phase of the TuPro clinical trial. In addition, the case study presented in this report demonstrated the ability of both methods to provide interpretable single-cell measurements useful for molecular tumor boards, which goes well beyond what is provided by routine pathology analysis. In particular, single cell measurements generated both by CyTOF and IMC provided a deep understanding of the heterogeneity of the tumor compartment including the identification of key pathways modulated in the different tumor cell types as well as their spatial organization. This led to the identification of potential resistance mechanism based on NGFR expression as well as upregulation of CyclinD1 downstream protein CDK6, which could not be apprehended at this resolution by bulk NGS analysis. In parallel, the precise immune cell profiling pointed towards potential benefits of treatments with novel immune checkpoints inhibitors. The generation of similar insights by routine pathology could only be done in an iterative process, which would be much more time consuming and would not allow for co-expression profile identification. Altogether, IMC and CyTOF single cell readouts could provide clues related to drug resistance mechanisms and alternative treatment options for this patient based on both the tumor and the immune cell characterization. A deeper understanding of resistance mechanisms is of pivotal importance for treatment guidance, hence helping future patient care.

CyTOF and IMC have both a strong potential to bring added value to the treatment decision making in the clinic. The validation and standardization strategies described for these cutting-edge technologies are crucial to ensure data quality required to enable their use in translational studies and eventually their integration in the clinical decision process made by oncologists.

## Acknowledgments

The authors acknowledge the support from the Biobank Team of the Department of Dermatology at the University Hospital Zurich as well as the Biobank Team of the Institute of Pathology and Molecular Pathology at the University Hospital Zurich who provided us with the samples used in this study. BB was supported by a SNF project grant and by the European Research Council (ERC) under the European Union’s Horizon 2020 Program under the ERC Grant Agreement no. 866074 (“Precision Motifs”).

## Author contributions

S.C., R.C., and B.B. conceived the study. S.S. performed the CyTOF experiments with the help of A.J., and S.C. S.E. performed the IMC experiments with the help of R.C. R.C. and S.C. performed the data analysis with the support of S.X. M.P.L. and R.D. contributed to the interpretation of the clinical data. R.C., S.C., and B.B. wrote the manuscript with input from all authors.

## Declaration of Interests

The authors declare no competing interests concerning the data presented in this manuscript.

---

**Figure S1. Analyses of references and of effect of freezing on antibody panel staining efficiency, related to Figure 1**. A-B. t-SNE representations of a maximum of 3,000 cells from each reference used as controls for A) the CyTOF immune panel and B) the IMC panel. C-D. Heatmap of mean scaled counts for each marker in each reference sample stained with C) the CyTOF immune panel and D) the IMC panel. E-F. t-SNE representations of E) a maximum of 3,000 cells from each reference stained with the fresh and the frozen antibody panels for CyTOF immune panel and F) all reference cells for IMC panel. Cells on the t-SNE are colored by antibody panel type. G. Density plot for each individual marker included in the CyTOF tumor panel across all the reference samples included in the study. For markers indicated with asterisks, only the most positive reference was used. The data obtained with the fresh and the frozen antibody panels are overlaid in each plot. H-I. Dot plots of median signal intensities in H) the reference sample with the highest expression level for each marker upon staining with fresh or frozen CyTOF immune panels and I) all pooled reference cells for the IMC analysis. The markers in each pair for fresh vs. frozen are connected with a line.

**Figure S2. Data stability achieved over a year of acquisition for the CyTOF immune panel, related to Figure 2**. A-B. Stacked bar plots of A) tumor-immune frequencies and B) all immune subsets identified in each experiment with reference samples acquired with the CyTOF immune panel. C. Dot plots of CVs of the frequencies of immune subsets. Dashed lines indicate CVs of 15% and 30%. D. Dot plots of median signal intensities of indicated markers in the reference samples across all measured time points in CyTOF experiments with the immune panel. Marker expressions in each cell type across experiments are connected with lines. E-F. t-SNE representations of E) a maximum of 2,000 cells from each of the 12 identified immune populations and F) corresponding immune markers in granulocyte populations. G. Dot plots showing the median signal intensities of markers in the reference samples across all measured time points using IMC. Marker expressions in each cell type across experiments are connected by lines.

**Figure S3. Batch correction upon linear scaling based on a percentile, related to Figure 3**. A-C. Density plots of all markers included in A) the CyTOF tumor panel, B) CyTOF immune panel, and C) IMC panel before (top) and after (bottom) batch correction.

**Figure S4. Validation of batch correction on independent tumor samples, related to Figure 4**. A. t-SNE representations of a maximum of 1,000 cells from the validation tumor samples for each acquisition batch colored by scaled intensities for each marker. B. Boxplots of CVs for all markers above the threshold of 0.15 mean scaled intensity (raw, batch corrected) for the three most abundant tumor clusters.

**Figure S5. Single-cell readouts of the tumor compartment of a metastatic melanoma cancer patient sample for clinical decision support, related to Figure 5**. A. Dot plot of numbers of cells from the sample from the metastatic melanoma cancer patient (red dots) in the indicated analyses, overlayed on the boxplot corresponding to the data collected for the entire cohort B. UMAP representation of all tumor cells from the patient colored by FlowSOM cluster. C. The same UMAP representation, colored by scaled intensities for melanoma markers. D. IMC images from three central regions of the patient sample showing the expression of MelanA (blue) NGFR (green), and GLUT1 (red) (top), and corresponding single-cell data showing the identified tumor clusters (bottom). Scale bar, 250 µm. E. Stacked bar plot of frequencies of identified tumor clusters by imaged regions. F. Dot plot of the Clark-Evans scores for imaged regions. The color scale and dot size correspond to scores and p-values, respectively.

**Figure S6. Single-cell readouts of the immune compartment of a metastatic melanoma cancer patient sample for clinical decision support, related to** Figure 6. A. t-SNE representation of a maximum of 1,000 immune cells identified by CyTOF in the melanoma patient sample colored by scaled intensities of immune markers. B. UMAP representation of a maximum of 20,000 immune cells identified by IMC colored by scaled intensities of immune markers. C. Boxplots showing the distributions of cell frequencies identified by IMC across the TuPro cohort. Red dots indicate the values for the analyzed patient sample.

## Methods

### Clinical samples

Melanoma tumor samples were collected at the University Hospital Zurich, in the context of the Tumor Profiler clinical study (Registration IDs: 2018-02050 (KEK ZH, Switzerland), 2018-02052 (EKNZ, Basel, Switzerland), 2019-01326 (KEK ZH, Switzerland)). For IMC analysis, melanoma tissue sections, the reference cell pellets as well as the reference tonsil tissue were prepared from FFPE tissue blocks by the Department of Pathology and Molecular Pathology, University Hospital Zürich. Both tumor samples and reference tissues were cut to a thickness of 3 µm and placed on the same slide before staining and acquisition. For CyTOF analysis, material remaining after pathological assessment were prepared as slow frozen material by the SKINTEGRITY.CH Biobank (Dermatology Department of the University Hospital Zürich) as recently described (37).

### Cell lines

Human cell lines, including HMLE, LNCaP, PC-3, A375, G361, KURAMOCHI, OVCAR3, SKOV-3, and A431 were obtained from the American Type Culture Collection (ATCC) and cultured according to ATCC recommendations. MKFY6, MNAT5, and MTG5K lines are early passage cells derived from melanoma patients prepared by the Dermatology Biobank (Department of Dermatology at University Hospital Zurich). PBMCs from healthy donors were obtained from the Zurich Blood Transfusion Service and were isolated by histopaque (Sigma Aldrich) density gradient centrifugation. PBMCs were stimulated with either 0.1 μg/mL phytohemagglutinin for 24 h or 1 μg/mL lipopolysaccharide for 48 h.

### Tissue preparation for CyTOF

Slow frozen tissue samples were kept at -80 °C until dissociation. At this stage, frozen tissue pieces were quickly thawed at 37 °C, rinsed with RPMI, and minced into 0.5-1 mm pieces. Enzymatic dissociation was performed in presence of collagenase (Worthington), DNase I Type 4 (Sigma), accutase (Sigma), and hyaluronidase (Sigma) for 30 min at 37 °C under gentle agitation on the MACSmix (Miltenyi Biotec). The resulting single-cell suspension was filtered sequentially through sterile 70-μm and 40-μm cell strainers. Cell purity was assessed using the NucleoCounter (Chemometec), and when the percentage of debris was above 50%, a debris removal step was performed using the Debris Removal Solution (Miltenyi Biotec) using the manufacturer’s procedure. If cell viability was below 50% after debris removal, dead cell exclusion was performed using an EasySep™ separation kit (Stem Cell Technologies) according to the manufacturer’s instructions. The cell suspension was stained for viability with 25 μM cisplatin (Enzo Life Sciences) in a 1-min pulse before quenching with 10% FBS. Cells were then fixed with 1.6% paraformaldehyde (Electron Microscopy Sciences) for 10 min at room temperature and stored at -80 °C.

### Mass cytometry barcoding

Before staining, cells were barcoded using a 20-well barcoding scheme consisting of unique combinations of three out of six barcoding reagents as previously described using palladium isotopes (^102^Pd, ^104^Pd, ^105^Pd, ^106^Pd, ^108^Pd, and ^110^Pd, Standard BioTools) chelated to 1-(4-isothiocyanatobenzyl) ethylenediamine-N,N,N’,N’ tetraacetic acid (Dojino). References were barcoded in position 1 to 10 as a master mix as previously described (2), aliquoted, and frozen at -80 °C as a dry pellet. During each experiment, samples from up to five patients (blood and tumor tissue) were barcoded and spiked in the reference set. Pooled cells were split in two and stained with the immune and the tumor panels.

### Antibodies and antibody labeling

The antibodies used in this study, including provider, clone, and metal tag, are listed in Table S1. Antibody conjugation was performed using the MaxPAR antibody labeling kit (Standard BioTools) according to the manufacturer’s procedure. Upon conjugation, the yield of recovered antibody was assessed on a Nanodrop (Thermo Scientific) and then diluted with Candor Antibody Stabilizer (by at least 50%). We performed titrations to determine optimal concentrations of all conjugated antibodies, which were managed using the cloud-based platform AirLab (38).

### Sample staining and CyTOF data acquisition

For CyTOF experiments, cells were incubated with FcR blocking reagent (Miltenyi Biotec) for 10 min at 4 °C and were subsequently stained with 100 µL of the antibody panel per 10^7^ cells for 45 min at 4 °C. Cells were washed three times in cell staining media (CSM, PBS 0.5% BSA and 2mM EDTA), once in PBS, resuspended in 0.4 ml of 0.5 µM nucleic acid Ir-labeled intercalator (Standard BioTools) and incubated overnight at 4 °C. Samples were then prepared for CyTOF acquisition by washing the cells once in CSM, once in PBS, and once in water. Cells were then diluted to 0.5 x 10^6^ cells/mL in Cell Acquisition Solution (Standard BioTools) containing 10% EQ™ Four Element Calibration Beads (Standard BioTools). Samples were acquired on a Helios upgraded CyTOF 2 in independent FCS files.

### CyTOF data preprocessing and automated cell classification

Individual FCS files collected from each set of samples were pre-processed using a semi-automated R pipeline based on CATALYST to perform individual file concatenation, bead-based normalization, compensation, debarcoding, and batch correction as previously described (25). During this process, inter-sample doublets were excluded based on the debarcoding scheme and intra-sample doublets were excluded based on DNA content. Spillover matrix for CyTOF compensation was assessed on all antibodies used in this study as previously described (39). After pre-processing, a subset of 1,000 randomly selected cells from each sample were exported as FCS files and loaded into Cytobank. Immune cell subsets were manually gated according to the profiles shown in Figure S1C. FCS files corresponding to each gate were exported and used to train a random forest classifier (R package randomForest) (40) based on 500 trees and 6 variables tried at each split, leading to an out-of-bag estimate of error of 0.43%. The resulting random forest model was used to assign each cell of the dataset to the predefined cell types. Based on a 40% assignment probability cutoff and a 20% delta cutoff between the first and the second assignment, 98% of the cells were retained in the analysis. Tumor clusters were identified by unsupervised clustering based on self-organizing maps (R package FlowSOM) (41).

### IMC reference sample preparation

The IMC reference samples, except for the tonsil tissue, were assembled using PBMCs, melanoma cell lines (A375, G361, MKF6Y, MNAT5, MTG5K), and ovarian cancer cell lines (KURAMOCHI, SKOV3, OVCAR3). Between 25 and 35 million cells were harvested in suspension and centrifuged for 5 min at 250 *g*. Cells were washed three times by removing the supernatant and resuspending in PBS. A final supernatant removal was performed before adding plasma followed by stirring until the formation of a single cell suspension. Thrombin was then added, quickly mixed, and incubated for 5-10 min until a clot was formed. The pellets were transferred to a biopsy capsule with a small mesh and then into a larger capsule. Cell pellets were fixed using 10% buffered formalin for 12-24 h, washed twice in PBS for 15 min to remove excess formalin. Cell pellets were finally washed using 50% EtOH for 30 min and were stored in 70% EtOH at 4 °C until embedding.

### IMC sample staining and data acquisition

Sample were dewaxed using the AS-2 Automatic staining system (Pathisto GmbH) with the following program: 3x10 min Ultraclear, 2x 5 min 100% EtOH, 2x 3 min 96% EtOH, 2x 3 min 90% EtOH, 1x 3 min 80% EtOH, 1x 3 min 70% EtOH, and Tris-buffered saline (TBS) for 5 min. This was followed by a 30-min antigen retrieval using Tris-EDTA pH 9.2 at 95 °C in a Decloaking chamber (NXGen). Samples were cooled at room temperature in the antigen retrieval solution for 20 min and then washed for 10 min in TBS. Blocking with 3% BSA in TBS with 0.1% Tween was performed for 1 h at room temperature. The antibody panel was mixed in a blocking buffer and incubated with the sample overnight at 4 °C. Samples were warmed to room temperature and stained with iridium (500 µM 1:100 in TBS) for 7 min, followed by three 5-min washes with TBS. After a short dip into doubly distilled H_2_O, samples were dried with compressed air.

### IMC image acquisition and processing

IMC images were acquired using a Hyperion Imaging System (Standard BioTools). Each reference sample was ablated with a laser frequency of 200 Hz. Raw image data were converted into image stacks as described in the ImcSegmentationPipeline (https://zenodo.org/record/3841961#.Y1vp5sFByFo). A customized U-Net model (42) was used to predict cell compartments (nuclei, cytoplasm) as a basis for cell single segmentation. To perform this task, specific structural markers were selected for their ability to identify nuclei and cytoplasm before performing cell segmentation. The resulting maps were segmented using profiler and cell marker intensities were also extracted using CellProfiler after spillover compensation. Single-cell data from each reference sample were exported as FCS files and loaded into Cytobank. The main cell populations were annotated using gates based on their marker expression as described in Figure S1D. The resulting annotated cells were imported to R and used to train a random forest classifier (R package randomForest) (40) based on 500 trees and 6 variables tried at each split, leading to an out-of-bag estimate of error of 0.2%. The model was used to predict each cell of the dataset. Cell populations from the metastatic melanoma cancer patient were annotated with a combination of unsupervised clustering using the Leiden algorithm (R package igraph) (43) and a random forest classifier (R package randomForest) (40). The tumor clusters were identified by unsupervised clustering based on self-organizing maps (41). The neighborhood permutation testing (28) was performed after constructing a cell neighborhood graph of cells within a distance of 20 µm between each other (R package imcRtools, method=classic, n=999 permutations). Clark-Evans indexes were computed using spatstat, a dedicated R package for point pattern analysis (44).

## Notes

### Competing Interest Statement

The authors have declared no competing interest.

### Summary of Updates

Figures size updated. Updated labels for Figure 6J.

