## Supplementary figures and images for "Standardization of suspension and imaging mass cytometry readouts for clinical decision making"

### Supplemental Figure 1

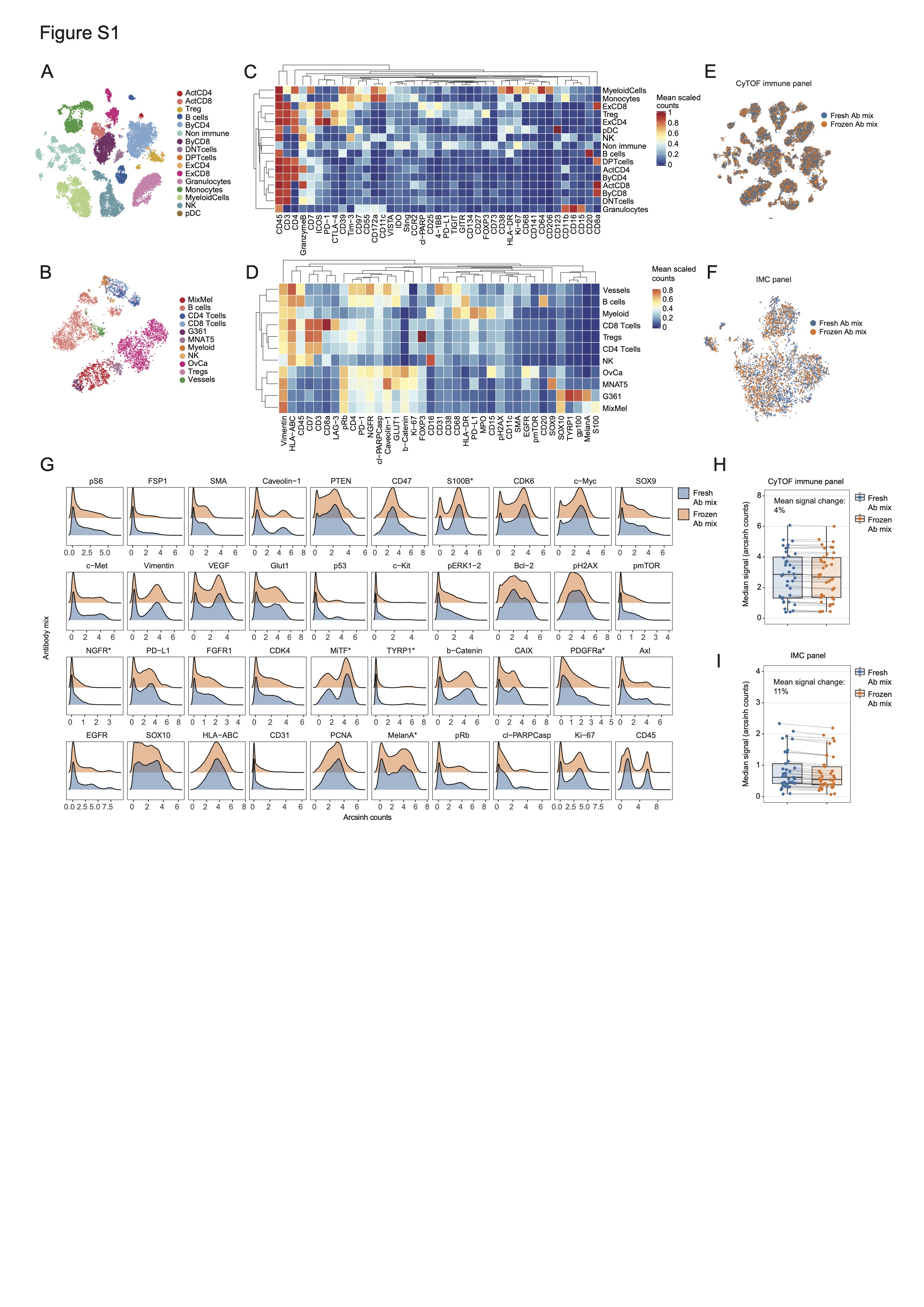

### Supplemental Figure 2

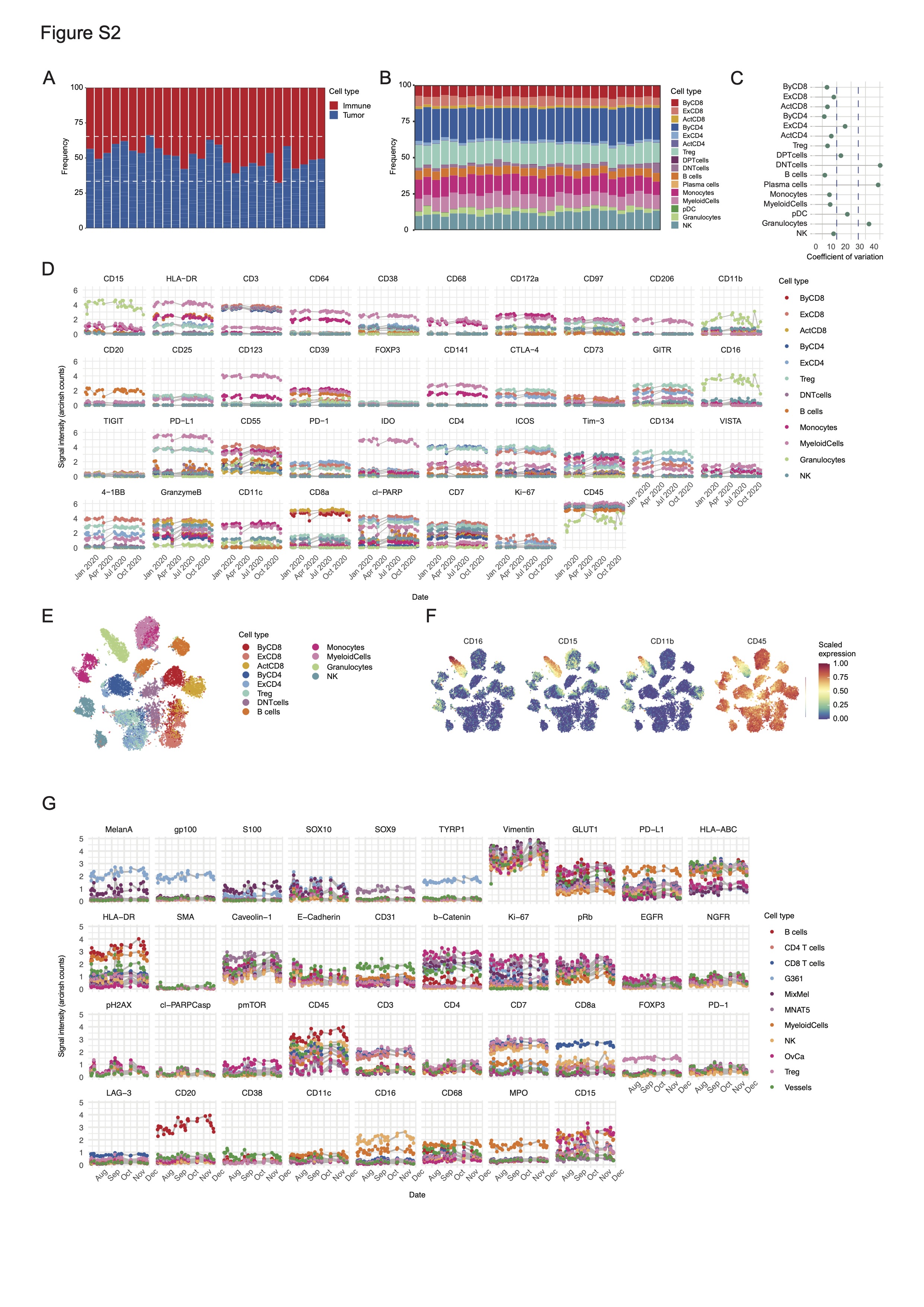

### Supplemental Figure 3

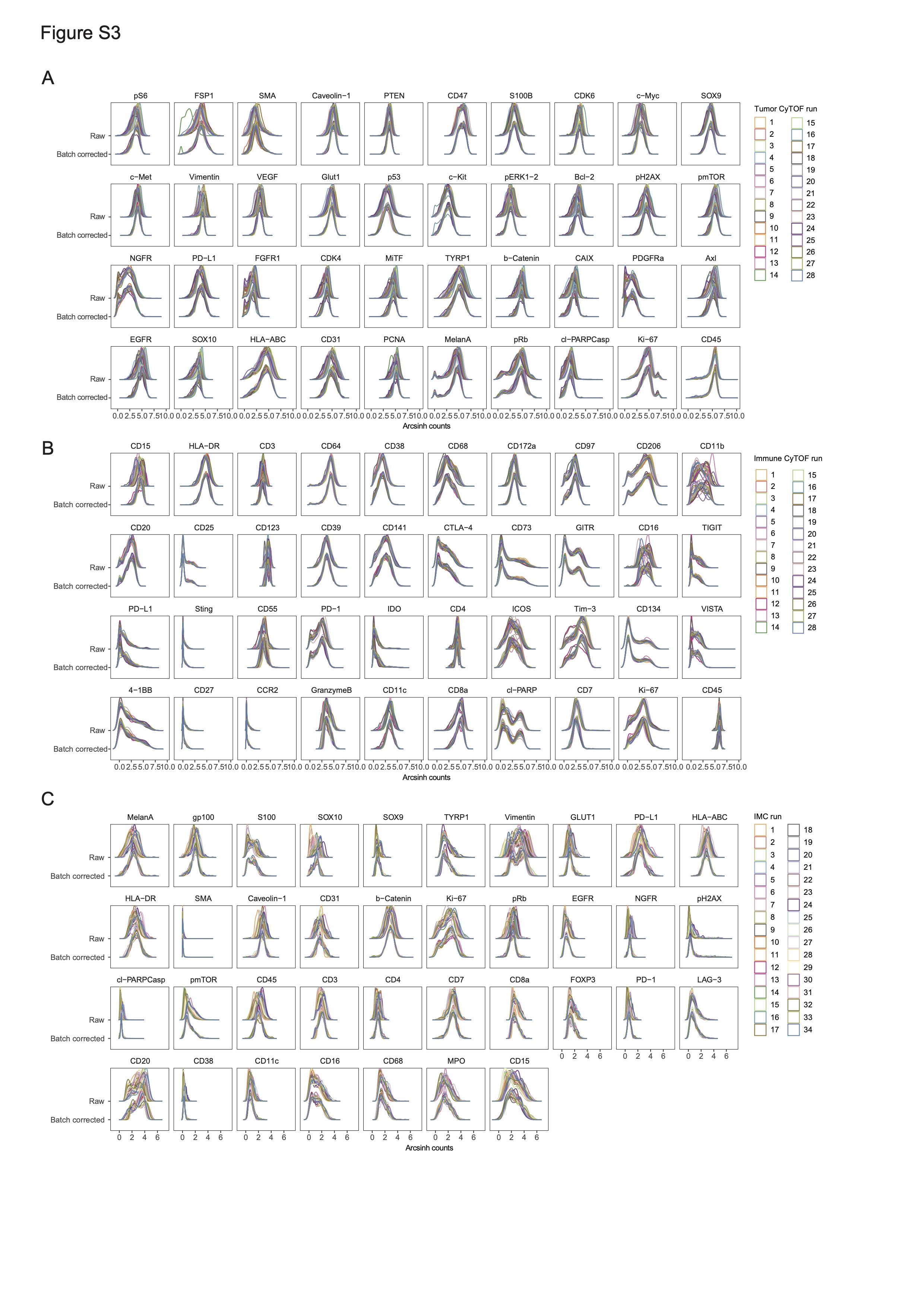

### Supplemental Figure 4

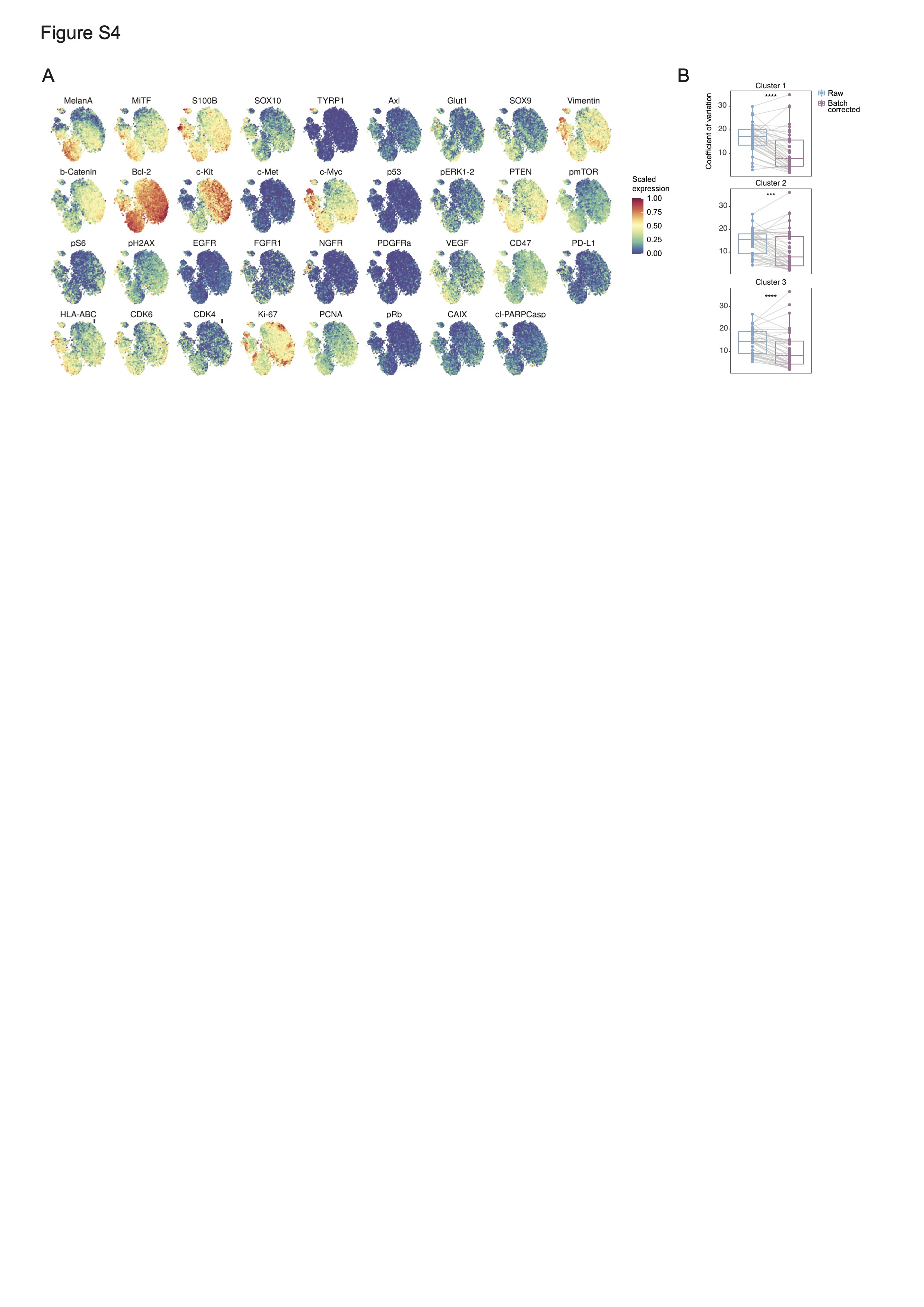

### Supplemental Figure 5

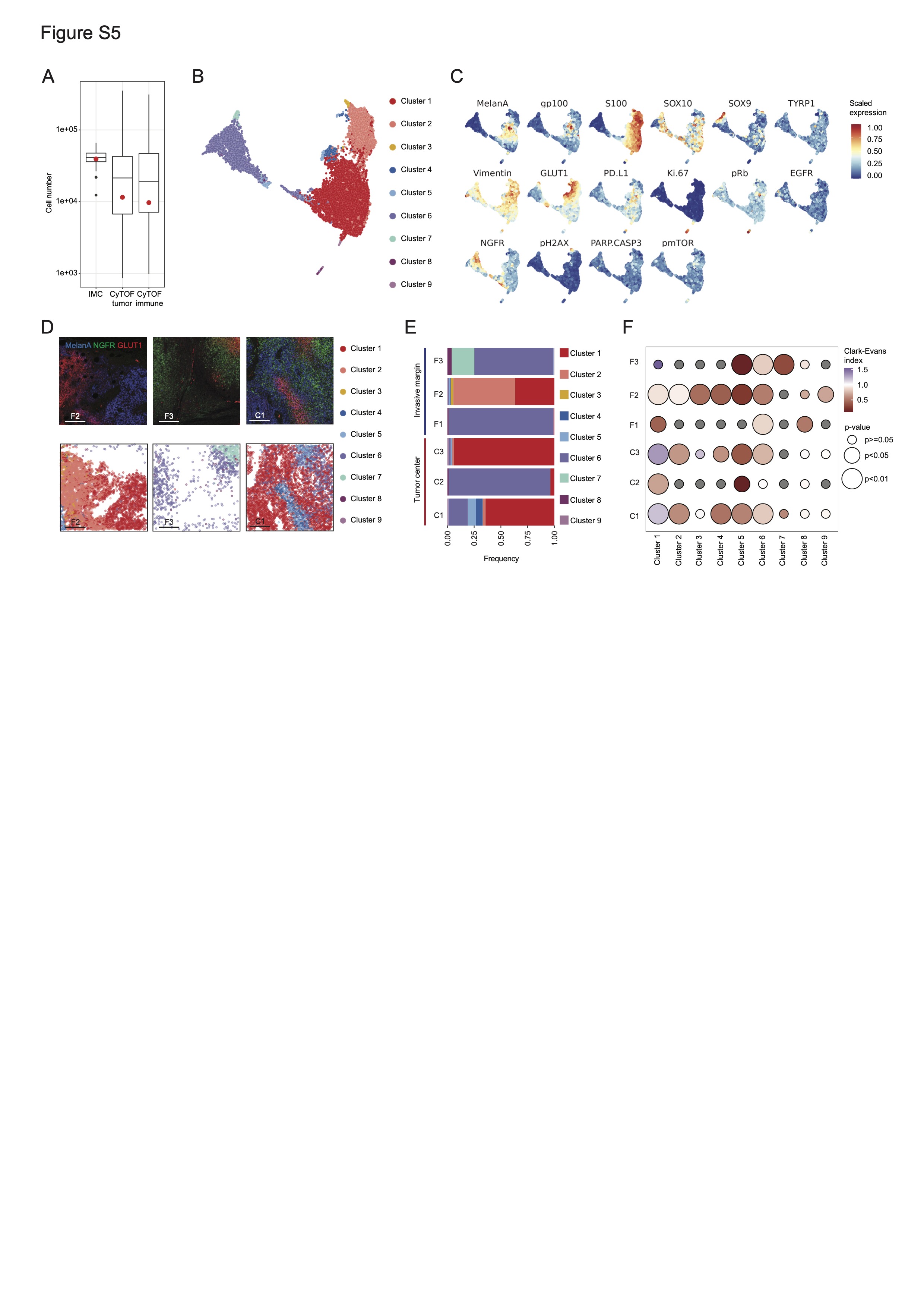

### Supplemental Figure 6

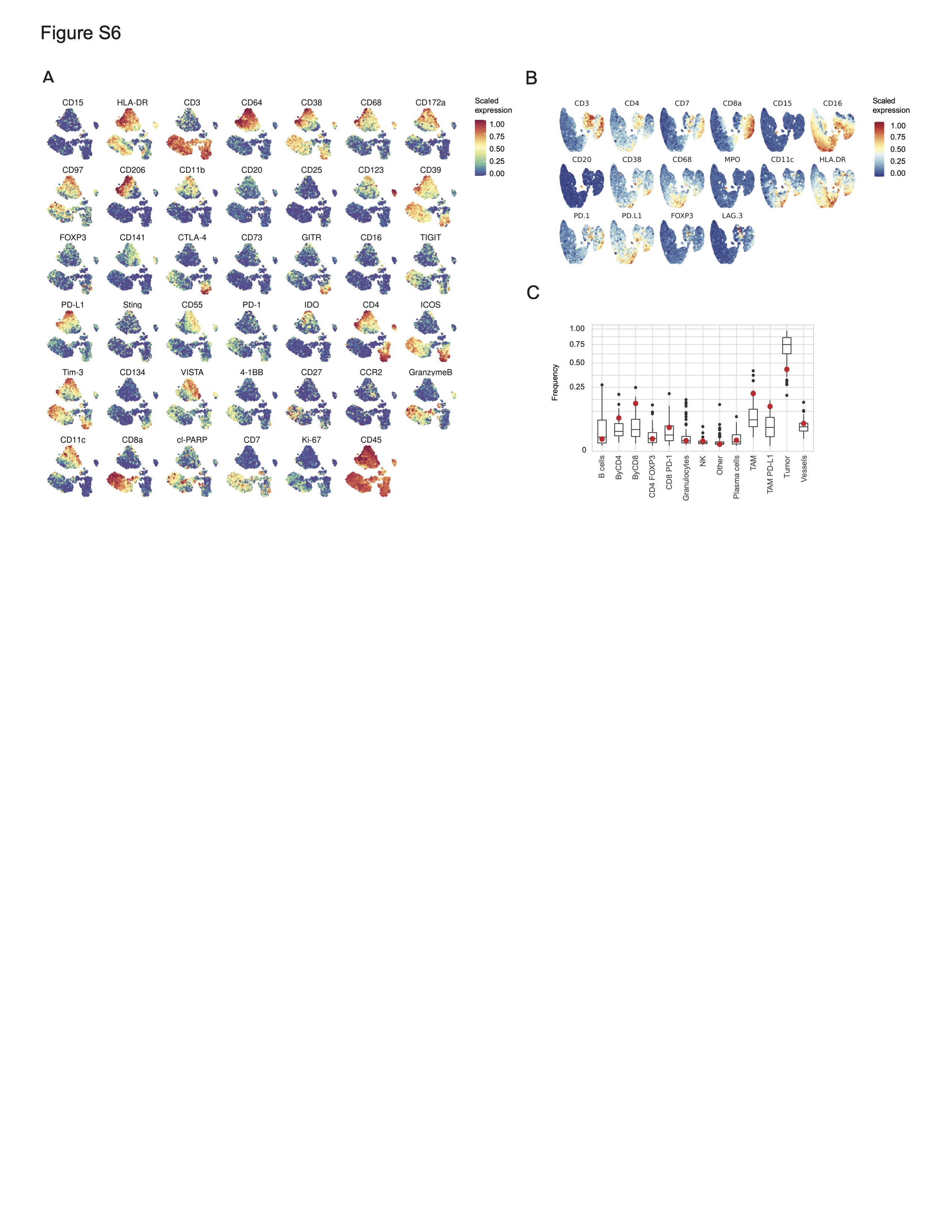
